# An enzyme-based system for the extraction of small extracellular vesicles from plants

**DOI:** 10.1101/2021.12.22.473784

**Authors:** Qing Zhao, Guilong Liu, Manlin Xie, Yanfang Zou, Zhaodi Guo, Fubin Liu, Jiaming Dong, Jiali Ye, Yue Cao, Ge Sun, Lei Zheng, Kewei Zhao

## Abstract

Plant-derived nanovesicles (NVs) and extracellular vesicles (EVs) are considered to be the next generation of nanocarrier platforms for biotherapeutics and drug delivery. However, EVs exist not only in the extracellular space, but also within the cell wall. Due to the limitation of isolation methods, the extraction efficiency is low, resulting in the waste of a large number of plants, especially rare and expensive medicinal plants.There are few studies comparing EVs and NVs. To overcome these challenges, we proposed and validated a novel method for the isolation of plant EVs by degrading the plant cell wall with enzymes to release the EVs in the cell wall, making it easier for EVs to break the cell wall barrier and be collected. We extracted EVs from the roots of Morinda officinalis by enzymatic degradation(MOEVs) and nanoparticles by grinding method (MONVs) as a comparison group. The results showed smaller diameter and higher yield of MOEVs.Both MOEVs and MONVs were readily absorbed by endothelial cells without cytotoxicity and promoted the expression of miR-155. The difference is that the promotion of miR-155 by MOEVs is dose-effective. More importantly, MOEVs and MONVs are naturally characterized by bone enrichment. These results support that EVs in plants can be efficiently extracted by enzymatic cell wall digestion and also confirm the potential of MOEVs as therapeutic agents and drug carriers.

## 1 Introduction

Currently, several researchers have developed nanovesicles (NVs) from plants. These NVs have a structural composition similar to that of mammalian exosomes[1] and can regulate biological functions across borders in animals and humans[2–6]. Several recent studies have shown that plant-derived NVs have intrinsic therapeutic activities, such as maintaining intestinal stem cells and shaping the intestinal microbiota to enhance intestinal barrier function to relieve colitis[7], improving anti-inflammatory properties in intestinal diseases[8], preventing alcohol-induced liver injury [9], promoting wound healing[10], and participating in tumor immunomodulation[3]. In addition, these vesicles have drug-carrying properties.For example, Wang et al. Loaded methotrexate (MTX) into grapefruit-derived NVs and found that MTX had significantly reduced toxic effects and significantly improved the therapeutic effect of dextran sodium sulfate-induced colitis in mice [11]. Plant-derived NVs are considered to have potential as therapeutic agents or drug carriers due to their small size and low immunogenicity, as well as the advantages of a wide range of medicinal plant sources, not carrying human or zoonotic pathogens and unique therapeutic activity compared to mammals[12].

However, plants do not have organic liquids for direct extraction of EVs like mammals.It was found that EVs structures exist not only in the extracellular space, but apparently also within the cell wall[13].There are two main methods for the extraction of plant-derived NVs and EVs[14], the most commonly used method is grinding[8, 15, 16], although this method can prepare large amounts of NVs, however, since these vesicles are derived from whole plant tissues subjected to cellular destruction, this method is considered to extract mixed vesicles, not pure EVs.These vesicles are uneven in size and activity, which will bring certain influence to subsequent studies.The other method is apoplastic fluid extraction by vacuum osmosis[14]. Only a few EVs are extracted by this method, and due to the cell wall barrier effect, only a small number of EVs were allowed to pass through. Its low extraction efficiency resulted in the waste of a large amount of plant raw materials and could not meet the needs of mass preparation. Therefore, the development of a more efficient separation method has become an urgent need for the study of plant-derived EVs.

Thus, We proposed a new method for the separation of plant-derived EVs, strictly speaking, a pretreatment method for the separation of plant-derived EVs. We degraded plant cell wall with digestive enzymes targeted at the main components, which not only promoted the release of EVs in the cell wall, but also facilitated the EVs to break through the cell wall barrier and enter the solution more easily, improving the output of isolated EVs, and more importantly, reducing the contamination of intracellular components. This method has been validated in the extraction of EVs from Morinda officinalis(MO).

MO is one of the four famous medicines in south China. As a medicinal part of Traditional Chinese medicine, the root of MO has a variety of biological activities, including anti-osteoporosis[17–19], anti-inflammation[20–22] and antioxidant activity[23]. In this study, enzyme digestion and grinding method were used to isolate EVs from the root of MO.According to possible origin and composition, the EVs extracted by the enzyme digestion we called MO derived EVs (MOEVs), and the NVs extracted by the grinding process we called MO derived nanoparticles(MONVs). We compared MOEVs and MONVs from many aspects to further verify the effectiveness of enzymatic isolation of plant-derived EVs.

## 2 Materials and methods

### 2.1 Animals and cell lines

Wild-type (WT) C57BL/6 mice were purchased from Spelford (Beijing) Biotechnology Co. All animal experimental protocols were approved by the Experimental Animal Ethics Committee of the Ruiye Model Animal Center. Mouse brain-derived Endothelial cells.3(b.End3) and human umbilical vein endothelial cells (h-UVECs) were purchased from Shanghai Meiwan Biotechnology Co. Cells were cultured in DMEM high sugar medium with 10% fetal bovine serum, 100 U/mL penicillin and 100 mg/mL streptomycin, all purchased from Gibco. Cells were cultured at 37°C and 5% CO_2_.

### 2.2 Separation of MONVs and MOEVs

MO is a fresh product of Taoist medicinal herbs originating from its origin in Zhaoqing, Guangdong. As shown in Figure1, 200g of washed MO was squeezed and filtered to obtain juice, then subsequently centrifuged at 500g for 10min, 2000g for 20min, 5000g for 30min, 10000g for 60min in order to remove large particles of impurities, each centrifugation step was left for 5min. The collected supernatant was centrifuged at 100000g for 70min, resuspended in PBS and then precipitated by 0.22μm. The protein concentration of vesicles was determined using the BCA protein assay kit.

**Fig. 1:**
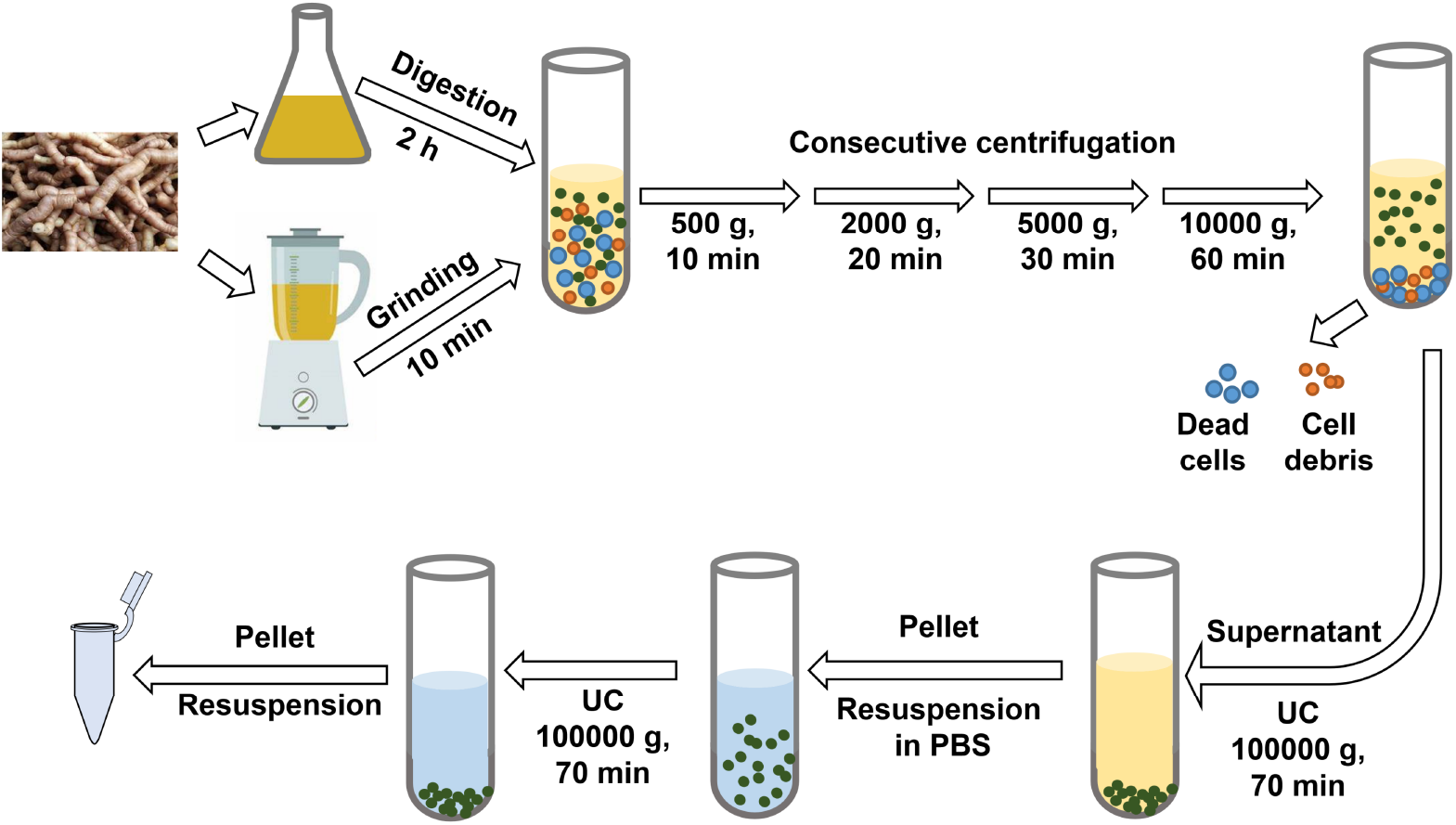
Scheme for isolation and preparation of MOEVs/MONVs

To isolate MOEVs, we referred to the method of preparing Protoplasts. In this experiment we aimed at enzymatic degradation of plant cell walls to allow easier release of extracellular vesicles into the solute, so we made necessary adjustments to the method of preparing plant protoplasts [24–27], which are briefly described as follows: vesicle isolation buffer (0.1% MES, 1 mmol/L CaCl2, pH 5.5), cellulase R10 and pectinase R10, hemicellulase (Cangzhou Xiasheng Enzyme Biotechnology Co.), and cell wall degradation was performed at 1:1:1 ratio and 0.5% concentration. The process was gently shaken and the juice was collected after 2 h. It should be noted that the fresh roots of MO were cut into 1 mm pieces before enzyme digestion to increase the surface area and accelerate the rate of enzyme degradation of the cell wall, and the necessary PBS washing step was required after cutting to remove intracellular vesicle contamination caused by broken cells, and the subsequent purification steps were the same as those for the juice extraction method, as shown in Figure1.

During isolation, samples were always kept on ice or stored at 4°C. After isolation, samples were stored at 4°C for less than 7 days or frozen at −80°C. Freeze-thaw was used no more than 1 time. Nano-Flow cytometric(NanoFCM) and transmission electron microscopy (TEM) assays were performed using fresh samples. For metabolomics analysis, all were used after freezing once in −80°C.

### 2.3 NanoFCM detection of diameters and concentrations of MONVs and MOEVs

Separated vesicles were assayed for diameter and concentration by nanoFCM, a technique for determining small particle size distribution profiles in suspensions using standards. Samples were diluted 1:100 and analyzed using a nano-flow cytometric analyzer (Xiamen-Flow N30) according to the manufacturer’s protocol. The basic procedure was as follows: a 250 nm QC particle calibration laser was used, which was used as a reference for particle concentration for the analysis. In addition, a mixture of particles of different sizes (68-155 nm) was used to determine the reference curve of the diameter distribution and the sample particle concentration and size distribution were calculated using NanoFCM software (NF Profession 1.0).

### 2.4 Transmission electron microscopy(TEM) detection of MONVs and MOEVs

5-10ul of sample solution was added dropwise to the copper grid. Sedimentation was performed for 3 min, filter paper was blotted from the edges to remove the floating solution, and phosphotungstic acid negative staining after PBS rinsing. The samples were dried at room temperature for 5 min and imaged on the machine using a TEM (JEM-1200EX, Japan Electronics Co., Ltd.) with an operating voltage of 80-120 kv for electron microscopy.

### 2.5 Flow cytometric viability assay

h-UVECs cells were cultured on proliferation medium. The h-UVECs cells were inoculated with 2000 cells/μl volume in 96-well plates and incubated for 12 h. The h-UVECs cells were treated with MONVs and MOEVs at particle concentrations of 0.5×10^9^ particles/ml and 4.5×10^9^particles/ml for 24 h and 48 h, respectively. h-UVECs cells were double fluorescently(Annexin V and PI) labeled and assayed for viability using a flow cytometer (Myriad).) to detect cell viability.

### 2.6 Cytotoxicity evaluation by CCK-8 assay

h-UVECs cells were cultured on proliferation medium. After incubation for 12 h, h-UVECs cells were treated with MONVs and MOEVs at 0.5×10^9^ particles/ml and 4.5×10^9^particles/ml concentrations for 12h, 24h, 48h and 72h, and the OD values at each time point were measured using an enzyme marker. Time point OD values using an ELISA.

### 2.7 Endothelial cell uptake assay

Vesicle entry was monitored by labeling MONVs and MOEVs with lipophilic DiI (Invitrogen) dye. After treatment of b.End3 and h-UVECs with labeled MONVs and MOEVs for 2h, 4h, and 8h.The proliferation medium was removed and the cells were fixed with 4% paraformaldehyde. Then Hoechst3342 (blue) was added to the cells and the nuclei were stained by incubation at room temperature for 15 min. Finally, the cells were washed with PBS containing 1% bovine serum albumin and the fluorescence (red and blue) was imaged under a fluorescence microscope (Leica). At least three fields of view were selected and analyzed using ImageJ software.

### 2.8 In vivo biodistribution assay

To analyze the biodistribution of MONVs and MOEVs in vivo, healthy female C57BL/6 mice (7-8 weeks) were injected intravenously with DiR-labeled MONVs and MOEVs. At the 12h, 24h and 48h after injection, the mice were executed and different organs were collected. DiR signal intensity from different organs was measured using an IVIS Lumina III in vivo imaging system (PerkinElmer) and analyzed with a Living Image in vivo imaging analysis system.

### 2.9 RNA extraction and determination of miR-155 expression in h-UVECs cells

To assess the miR-155 expression in endothelial cells, primers were designed as follows:

miR-155 (upper: 5’-TTAATGCTAATTGTGATAGGGGT-3’, lower: 5’-ACCCCTATCACAATTAGCAT-TAA-3’)

U6 (upper: 5’-ATTGGAACGATACAGAGAAGATT-3’, lower: 5’- GGAACGCTTCACGAATTTG-3’)

Total RNA was extracted from each group of cells using the TRIzol method, reverse transcribed into cDNA using the Biogenic miRNA First Strand cDNA Synthesis Kit, and cDNA of miR-155 and internal reference U6 was amplified using SYBR Premix Ex Taq II.

### 2.10 Metabolomics analysis

Equal amounts of each of all samples were mixed as quality control(QC) samples. The QC samples were interspersed between samples during the mass spectrometry loading process. Methanol, formic acid and acetonitrile were purchased from CNW, and L-2-chlorophenylalanine was purchased from Shanghai Hengchuang Biotechnology Co. All extraction reagents were pre-cooled at −20°C before use. The samples were first lyophilized, added with 400μL of methanol, vortexed for 30 s, and sonicated for 3 min. then two small steel balls were added and placed at −20°C for 2 min to pre-cool, and then grinded with a grinder (60 Hz) for 2 min. the grinded samples were centrifuged at 13000 rpm for 10 min at 4°C. 350μL of the samples were evaporated into the LC-MS injection vial, and the samples were extracted with The sample was re-dissolved with 300μL of aqueous methanol (V:V=1:4), vortex mixed for 30 s, sonicated for 3 min, and subsequently left to stand at −20°C for 2 h. The sample was centrifuged at 13000 rpm for 10 min at 4°C, and 100μL of the supernatant was aspirated with a syringe, filtered using a 0.22μm organic phase pinhole filter, and transferred to the LC injection vial for examination, and the analytical instrument for the experiment was The analytical instrument for the experiment was a Dionex U3000 UHPLC ultra-high performance liquid tandem QE high-resolution mass spectrometer consisting of a liquid-liquid mass spectrometer for LC-MS analysis. The chromatographic column: ACQUITY UPLC HSS T3 (100 mm×2.1 mm, 1.8 μm), 45°C, water (containing 0.1% formic acid) and acetonitrile (containing 0.1% formic acid) as mobile phases, flow rate 0.35 mL/min, injection 2μL. ion source: ESI, sample mass spectrometry signal acquisition using positive and negative ion scan mode, respectively.

The raw data were processed by metabolomics software Progenesis QI v2.3 software (Nonlinear Dynamics, Newcastle, UK) for baseline filtering, peak identification, integration, retention time correction, peak alignment and normalization. Compound identification was based on accurate mass numbers, secondary fragmentation, and isotopic distribution.Characterization was performed using The Human Metabolome Database (HMDB), Lipidmaps (v2.3), and METLIN databases, as well as self-built libraries. For the extracted data, the ion peaks with missing values (0 values) >50% within the group were removed and the 0 values were replaced by half of the minimum value, and the compounds obtained from the characterization were screened according to the compound characterization results scoring (Score) with a screening criterion of 36 out of 60, below 36 points were considered as inaccurate and deleted.

### 2.11 RNA, protein and lipid assay of MOEVs and MONVs

Lipids were purified from MOEVs and MONVs, chloroform: methanol: sample = 8:4:3 mixed and centrifuged at 10,000g for 10 min, divided into three layers (upper water middle protein layer), the bottom lipid layer was collected, dried at 100°C and resuspended with chloroform, and the lipids in the samples were detected by thin-layer liquid chromatography.

To determine whether MOEVs and MONVs contain RNA, extracted vesicles were aliquoted into two tubes each, added with QIAzol lysate, shaken vigorously and incubated for 30 min at room temperature, one tube with RNase A (Solarbio) and one tube without RNase A, incubated for 30 min at 37°C, and the RNA was extracted with the kit. 2.5% agarose electrophoresis was used to detect MOEVs and MONVs RNA.

Two vesicle suspensions of two concentrations were prepared in four separate tubes, two of which were stock solution and two of which were diluted 10-fold. Two of the tubes with different concentrations of vesicles were lysed by adding RIPA+PMSF (100:1, Solarbio, China) lysis solution, and the other two tubes without any reagent were used as controls. 10% SDS-PAGE gel was prepared according to the SDS-PAGE gel kit (Biyuntian, Shanghai, China). PAGE gels with a sample volume of 20ug per well, and after running the gels, staining was performed according to the instructions of the rapid silver staining kit (biosharp, China), and pictures were scanned using a scanner (Epson, Japan).

### 2.12 Differential metabolite pathway analysis

The pathway enrichment analysis of the differential metabolites helps to understand the mechanism of metabolic pathway changes in the differential samples. We performed metabolic pathway enrichment analysis of MONVs and MOEVs differential metabolites based on KEGG database (https://www.kegg.jp/).

### 2.13 Statistical analysis

Data obtained from the experiments are expressed as mean ± SD. statistical analyses were performed using oneway analysis of variance (ANOVA) and t-test test using GraphPad Prism (GraphPad Prism Software Inc., San Diego, CA, USA). Statistically significant at differences p<0.05, p<0.01 and p<0.001.

In the non-targeted metabolite analysis, unsupervised principal component analysis (PCA) was first used to observe the overall distribution between samples and the stability of the whole analysis process, and then supervised partial least squares analysis (PLS-DA) and orthogonal partial least squares analysis (OPLS-DA) were used to distinguish the overall differences in metabolic profiles between groups and to find the differential metabolites between groups.

In OPLS-DA analysis, Variable important in projection (VIP) can be used to measure the strength and explanatory power of the expression pattern of each metabolite on the categorical discrimination of each group of samples, and to explore the biologically significant differential metabolites. The t-test was further used to verify whether the differential metabolites between groups were significant. The screening criteria were VIP value >1 for the first principal component of the OPLS-DA model and p-value <0.05 for the t-test.

Correlation analysis was performed using Pearson correlation coefficient to measure the degree of linear correlation between two metabolites.

## 3 Results

### 3.1 Characterization and yields of MONVs and MOEVs

In order to compare the plant-derived EVs and NVs extracted by two different methods, we first analyzed the concentration and particle size of MONVs and MOEVs by NanoFCM. Most MONVs are distributed in the diameter range of 50-120nm, with an average peak diameter of 71.62 ± 2.296nm(n=9)(Figure2A and C). Most MOEVs are distributed in the diameter range of 50-80nm, with an average peak diameter of 65.46 ± 1.74nm(n=9)(Figure2B and C). MOEVs had a smaller size range and a more uniform diameter distribution compared with MONVs. TEM images also verified that both MONVs and MOEVs had typical exosome like morphology(Figure2D and E).The yield of MONVs and MOEVs was (1.41 ± 0.52) × 10^9^ particles and (4.35 ± 0.74) × 10^9^ particles per gram of tissue(Figure2 F). Protein concentration of MONVs (322.2 ± 10.51ug per gram of tissue)was also lower than that of MOEVs(423.8 ± 17.45ug per gram of tissue)(Figure2G).

**Fig. 2:**
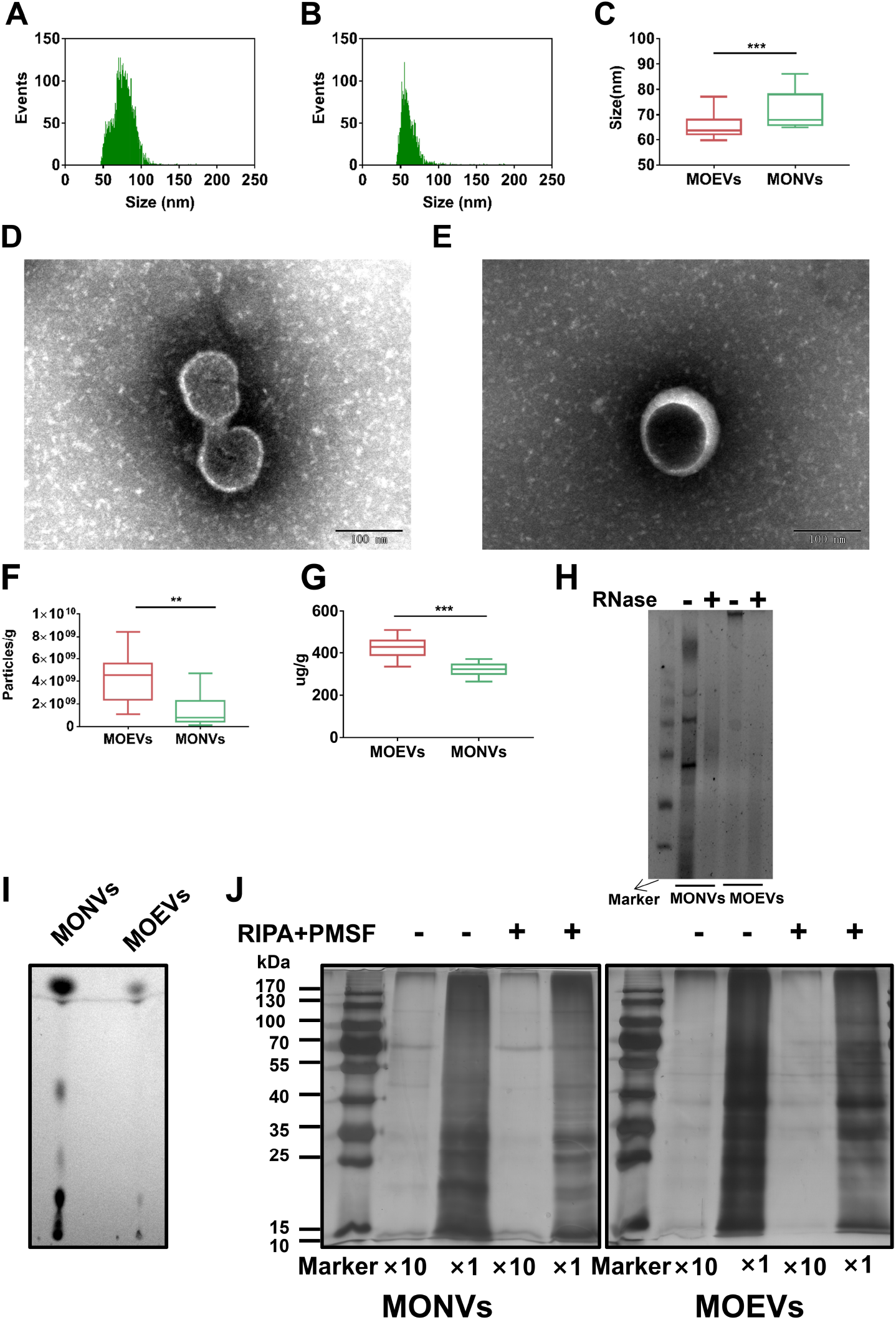
Characterization and yields of MONVs and MOEVs. (A,B) Diameter distribution of (A)MONVs(n=7103) and (B)MOEVs(n=3321) by nano-flow detection. (C) Diameter of MONVs and MOEVs. The average size potentials of MONVs and MOEVs were71.62 ± 2.296nm and 65.46 ± 1.74nm, respectively. (D,E) Morphology of (D)MONVs and (E) MOEVs analyzed by TEM. (F) Particle yields and (G) Protein yields of MONVs and MOEVs. (H) RNA gel electrophoresis,(I) Lipid and (J) Protein gel electrophoresis of MOEVs and MONVs.All values are expressed as mean ± SD (**p < 0.01, ***p < 0.001; n = 9)

In order to further understand the characteristics of the main chemical composition, lipids, RNA and proteins of MONVs and MOEVs were detected. Lipids and RNA were much higher in MONVs than in MOEVs, and RNA in MONVs and MOEVs was affected by RNA enzymes suggesting that these vesicles contain RNA (Figure2H and I).The total protein content and protein types in MOEVs were significantly higher than those in MONVs (Figure2J). These results suggest that the composition of MONVs and MOEVs differs, which may be related to their different origins. These results suggested that there were differences characteristics between the MONVs and MOEVs.

### 3.2 Effects of MONVs and MOEVs on efficiency of vascular endothelial cells(ECs) uptake

To determine whether MONVs and MOEVs have different effects on mammalian cell uptake, we compared the efficiency of vesicle uptake using the b.End3 and hUVECs. MONVs and MOEVs were labeled with DiI and then co-incubated with b.End3 and h-UVECs. We found that MONVs and MOEVs (red) were rapidly taken up by b.End3 and h-UVECs cells and preferentially localized in the cytoplasm of the cells (Figure3A and B). Flow cytometry shown that the percentage of h-UVECs cells that ingested DiI-MONVs and DiI-MOEVs increased with time and dose. The percentage of h-UVECs that ingested MONVs and MOEVs increased from 2.62% and 5.73% upto 24% and 35.6% for low concentrations compared to high concentrations at 2h, respectively. The percentage of h-UVECs cells ingesting MONVs and MOEVs increased from 4.78% and 11.20% to 55.4% and 75.7% for low concentrations compared to high concentrations at 8h. It showed that both MONVs and MOEVs were readily taken up by ECs, while the uptake efficiency of MOEVs was higher, which may be related to their smaller particle size (Figure3A).

**Fig. 3:**
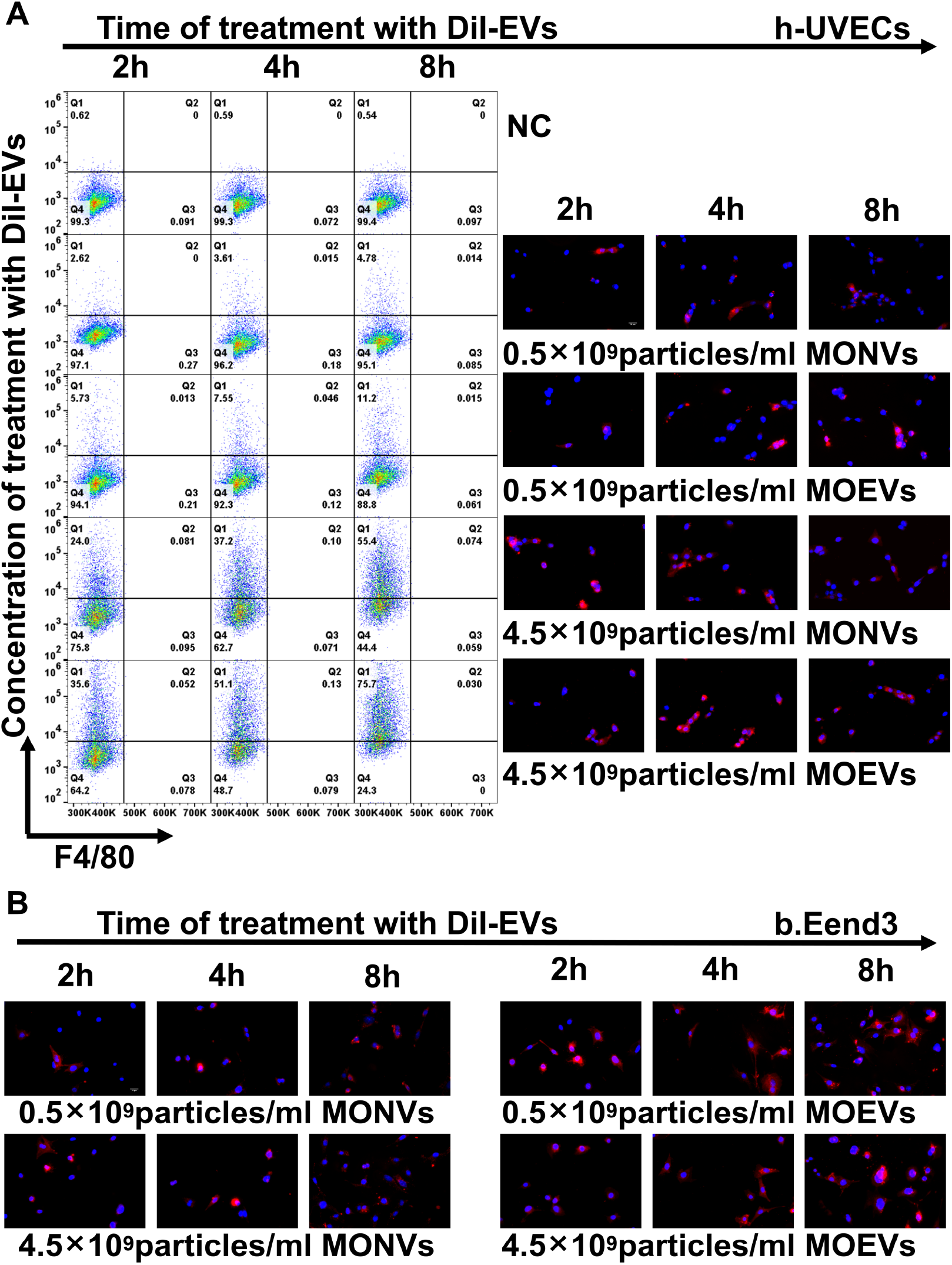
In vitro uptake of different concentrations of MONVs and MOEVs by b.End3 and h-UVECs at time profiles. (A) Dil-labeled concentration of different concentrations(0.5×10^9^particles/ml,0.5×10^9^particles/ml) MONVs and MOEVs were taken up h-UVECs and at (B)b.End3 different time(2h,4h,8h), respectively. MOEVs and MONVs were labeled by Dil in red, while the nuclei were labeled by Hoechst334 in blue.(scale bar:50μm, n = 3).

### 3.3 MONVs and MOEVs had no cytotoxicity and promoted the expression of miR-155 in endothelial cells(ECs)

The co-incubation of MONVs and MOEVs at different concentrations with h-UVECs showed no significant cytotoxicity within 72h. h-UVECs cells still maintained >90% viability at 24h and 48h co-incubation (Figure4A).Low concentration(0.5×10^9^/ mL)of MONVs and MOEVs significantly promoted the proliferation of h-UVECs at 48h and 72h, while high concentration (4.5×10^9^/ mL)of MONVs and MOEVs showed different performance(Figure4B). High concentration of MOEVs did not significantly promote the proliferation of h-UVECs, while high concentration of MONVs still promoted the proliferation of h-UVECs at 48h and 72h(Figure4B).These results suggest that both MONVs and MOEVs are not cytotoxic and also promote ECs proliferation.

**Fig. 4:**
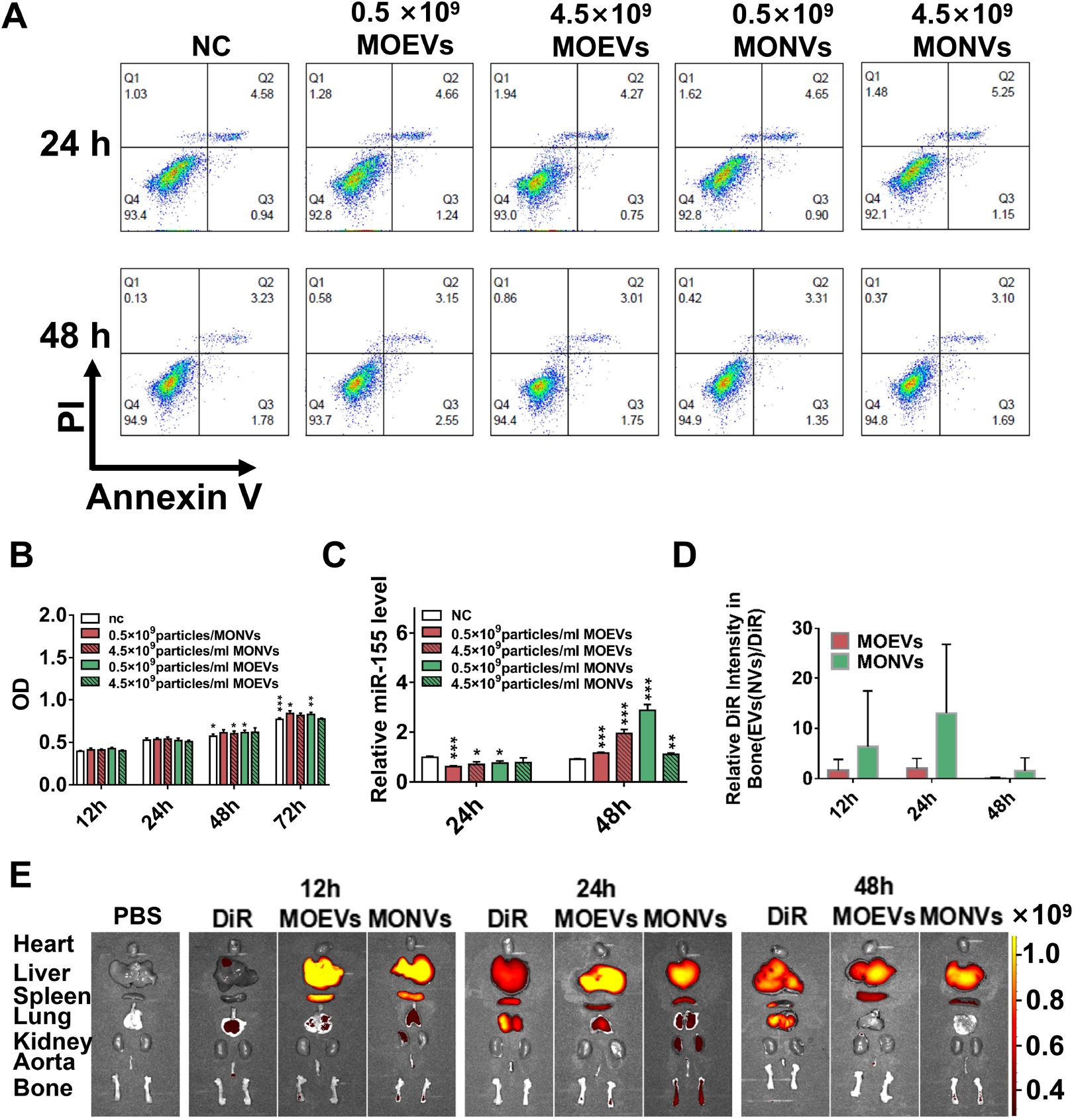
Effect of MOEVs and MONVs on the value-added, activity and promotion of miR-155 expression in h-UVECs cells. (A) Flow cytometry detection of the effect of different concentrations of MOEVs and MONVs on the activity of h-UVECs cells. (B) CCK8 assay of the effect of different concentrations of MOEVs and MONVs on the value-added of h-UVECs cells.(B) RT-PCR assay of histogram of the effect of different concentrations of MOEVs and MONVs on miR-155 expression in h-UVECs cells.In vivo organ distribution of MOEVs and MONVs (D) Relative fluorescence intensity histograms of bone tissue at 12h, 24h and 48h, with free DiR as the control group.(E) 12h, 24h and 48h in vivo organ distribution of MOEVs and MONVs mice.

To determine the effect of MONVs and MOEVs on the expression miR-155 in ECs,The expression levels of miR-155 in h-UVECs co-incubated with MONVs and MOEVs at 24h and 48 h were detected. MONVs and MOEVs both slightly inhibited the expression of miR-155 and were negatively correlated with the concentrations at 24h. MONVs and MOEVs strongly promoted the expression of miR-155 while only MOEVs was positively correlated with the concentrations at 48h (Figure4C).Generally speaking, MONVs and MOEVs have almost the same efficacy in regulating the expression of miR-155 in h-UVECs, but there are some differences mainly reflected in the concentration dependence of MOEVs at 48h.

### 3.4 Bone targeting ability of MONVs and MOEVs

Targeted therapy is one of the main methods to reduce adverse drug reactions.Therefore, we preliminarily evaluated the bone-targeting ability of MONVs and MOEVs. We observed strong fluorescence signal in both MONVs and MOEVs group mice at 12h and 24h.The fluorescence signal of MOEVs almost disappeared, while strong fluorescence signal could still be detected in the MONVs group at 48h. In general, both MONVs and MOEVs had bone targeting, and the fluorescence intensity changed with time(Figure4D and E).

### 3.5 Metabolomics of MONVs and MOEVs

The results of untargeted metabonomics showed regular and uncluttered basal peak pattern and the QC samples in PCA model diagram obtained through 7 cycles of cross verification were are closely clustered together, which proved that the instrument was stable and the data collected was reliable (Figure5A-C).17444 chromatographic peaks were extracted from MONVs and MOEVs, including 10051 positive ion mode(PIM) chromatographic peaks and 7393 negative ion mode(NIM) chromatographic peaks. Compared with database, 4018 compounds were identified, including 2704 in PIM and 1314 in NIM models(Figure5 D). Classification of 4018 identified compounds showed that MONVs and MOEVs showed similar species of compounds,and the main types include Lipids and functional-like molecules (29.76% vs 28.91%), Organic oxygen compounds (9.22% vs. 8.95%) and Organic acids and Derivatives (8.81% vs. 8.59%)(Figure5E). In addition, 27.35% and 26.15% of compounds in MONVs and MOEVs were not classified(Figure5E).We further analyzed the relative contents of the above compounds, and found that MONVs and MOEVs showed very different performances.The content of Lipids and lipid-like molecules in MOEVs was much higher than that in MONVs(87.62% vs. 66.17%; p<0.01)(Figure5F).

**Fig. 5:**
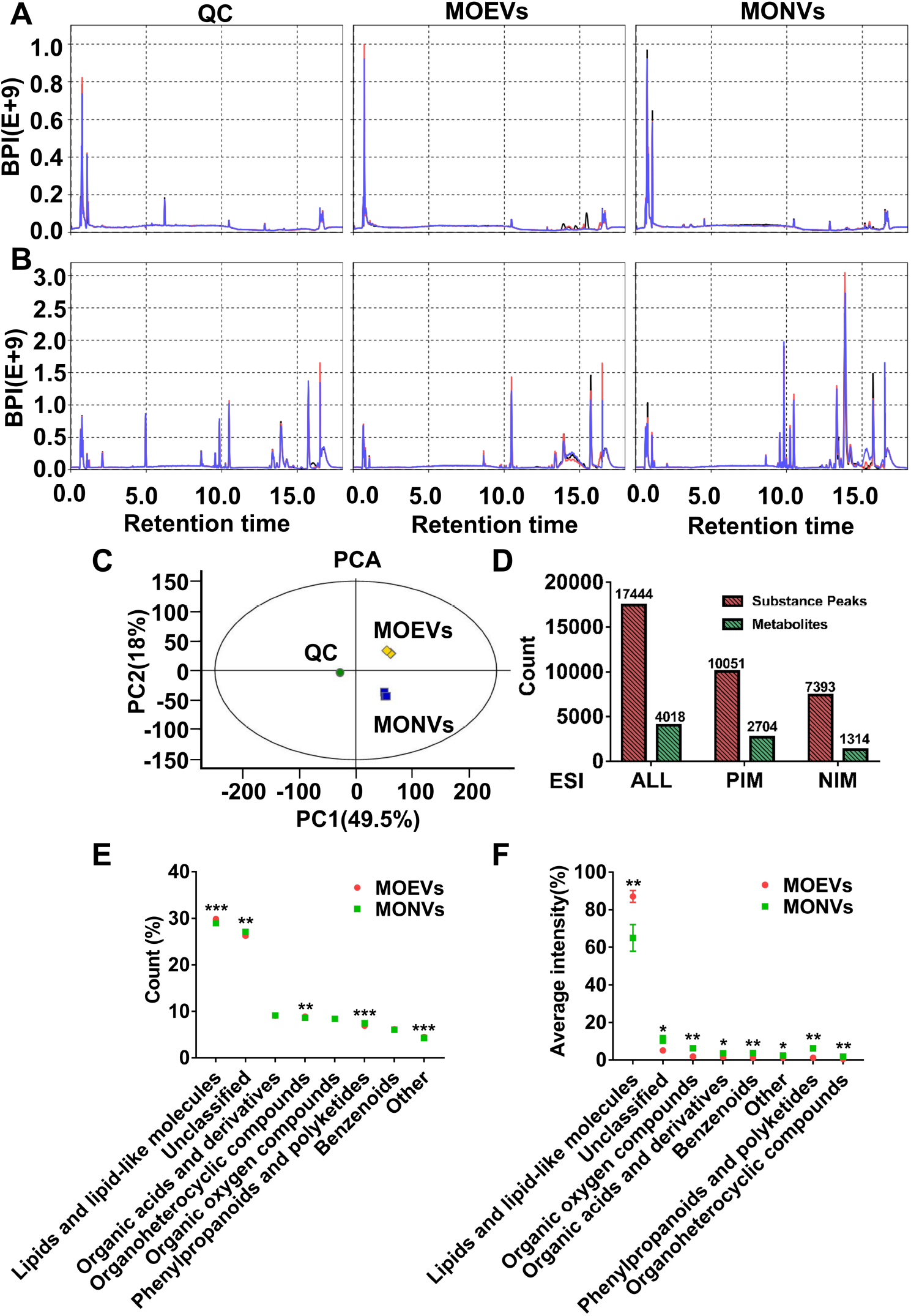
Quality control and substance confirmation of chromatographic-mass spectrometric detection of MOEVs and MONVs contents. (A) Base peaks of QC, MONVs and MOEVs in negetive ion mode and posittive ion mode. (C) Dot plots of PCA scores of QCs, MONVs and MOEVs. (D) Histogram of the number of peaks and substance confirmation for MONVs and MOEV in negetive ion mode(NIM) and posittive ion modes(PIM).(E,F)Composition of MONVs and MOEVs compounds.(E)Percentage of compound classification counts and (F)intensity.

Lipid composition was analyzed and the results showed that PC content was the highest (29.18%), followed by PE (14.86%) and PI (8.53%),in MONVs(Figure6A). PC accounted for 46.4% of lipids in MOEVs, 1.6 times higher than that in MONVs, followed by PE (29.06%) and PS (7.71%)(Figure 6B). Other lipids also had a large difference between MONVs and MOEVs (28.52% vs. 9.92%; p<0.01) (Figure 6A).Unsupervised principal component analysis (PCA) results showed R2X=0.806, indicating that MONVs and MOEVs could be well distinguished and the whole analysis process was stable and reliable (Figure6B).Supervised partial least squares (PLS-DA) results showed that MONVs and MOEVs could be completely separated, with model R2X(CUM)=0.907, R2Y(CUM)=1, and Q2(CUM)=0.995, indicating that the model had reliable stability and strong predictive ability(Figure 6C).Orthogonal partial least squares analysis (OPLS-DA) showed that MONVs and MOEVs had similar results (Figure6D). The OPLS-DA model was tested 200 times for response sequencing, and the corresponding OPLS-DA model was established to obtain R2=0.932 and Q2=0.1 of the random model, indicating that the model was of good quality (Figure6E).The load diagram visualized the intensity of the effect of metabolites on MONVs and MOEVs(Figure 6F).The Splot visualized the contribution rate of each variable to the grouping of MONVs and MOEVs (Figure 6G).The model parameters of the comparative analysis of the two groups were shown in Table 1.These analyses show that the model used was reliable and could be used to screen differential metabolites between MONVs and MOEVs.

**Table 1:**
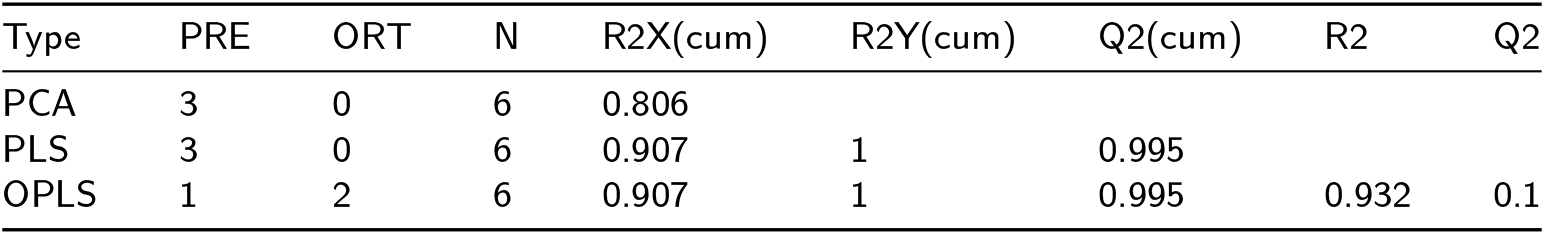
Table of model parameters for the comparative analysis of MOEVs and MONVs.

| Type | PRE | ORT | N | R2X(cum) | R2Y(cum) | Q2(cum) | R2 | Q2 |
| --- | --- | --- | --- | --- | --- | --- | --- | --- |
| PCA | 3 | 0 | 6 | 0.806 |  |  |  |  |
| PLS | 3 | 0 | 6 | 0.907 | 1 | 0.995 |  |  |
| OPLS | 1 | 2 | 6 | 0.907 | 1 | 0.995 | 0.932 | 0.1 |

**Fig. 6:**
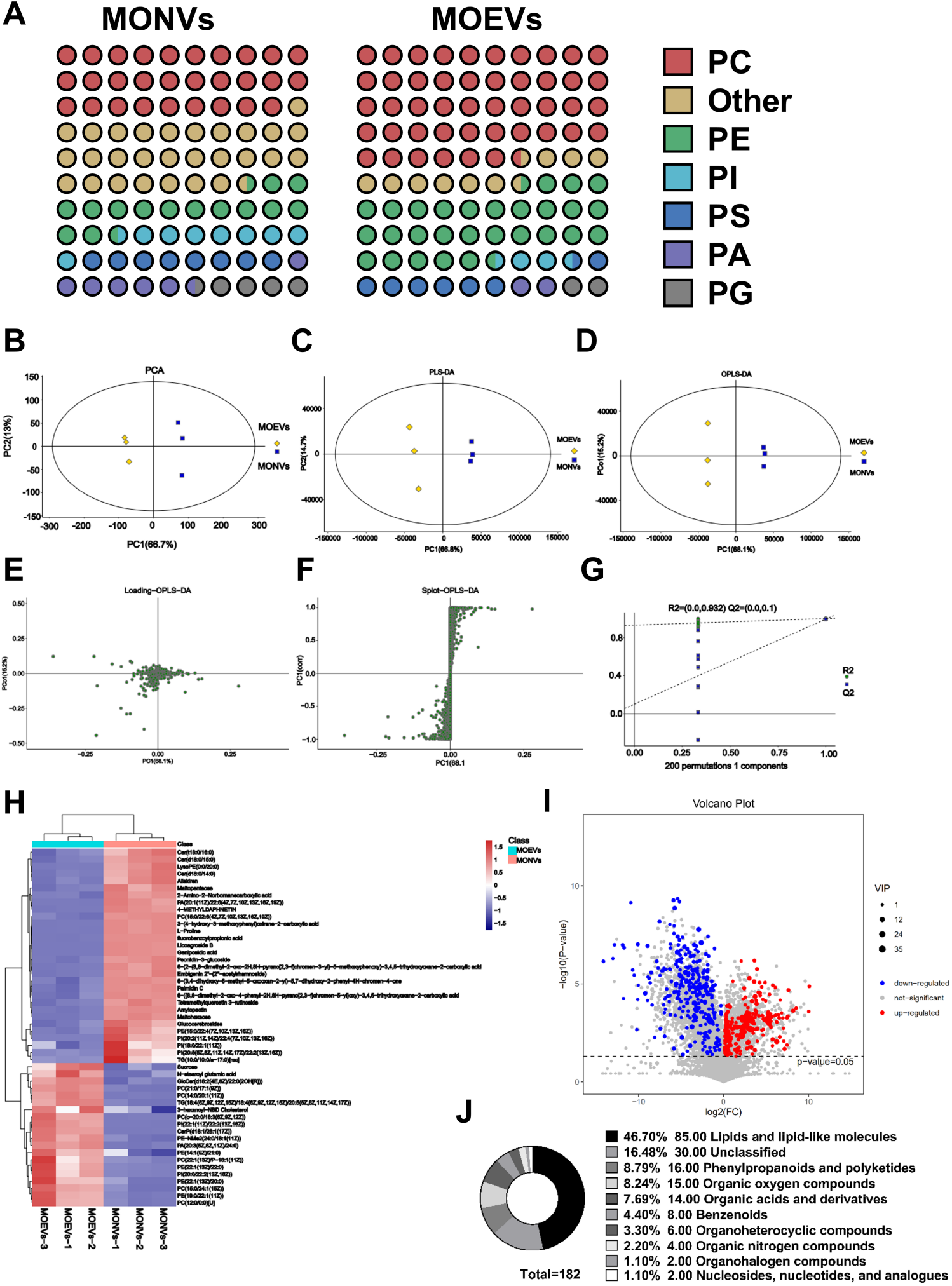
multivariate statistical analysis of lipid composition and compounds of MONVs and MOEVs. (A) Pie chart of lipid type content of MONVs and MOEVs. PC= phosphatidylcholine, PS=phosphatidylserine, PE=phosphatidylethanolamine, PI=phosphatidylinositol, PA=phosphatidic acid, PG=phosphatidylglycerol. (B) Plot of PCA scores of MONVs and MOEVs. (C) Plot of MONVs and MOEVs PLS-DA scores. (D) MONVs and MOEVs OPLS-DA score plots. (E) Permutation plot (F) Loading plot (G) Splot plot.Differential metabolite screening. (H) Heat map of TOP-50 differential metabolites.(I)Volcano plot of VIP and P screens for differential metabolites. (H) Heat map of TOP-50 differential metabolites.(J)Pie chart of Differential metabolite.

### 3.6 Differential metabolite analysis

The p value, VIP and Fold change values obtained by OPLS-DA model were visualized by volcanic map, which is conducive to screening differential metabolites.182 compounds with significant differences were screened according to preset rules. Compared with MONVs, 114 metabolites were down-regulated and 68 metabolites were up-regulated in MOEVs. The top five compounds with the highest VIP values, PE (19:0 / children of z) (11), PC (15:0/24:1 (z) 15), PC (o - 20:0 / and (z) 9 z, 6 z, 12), PE (children of 13 (z) / 22:0) and PE (children of 13 (z) / 20:0), were all lipids(Figure 6I). In order to display the relationship and the expression differences of metabolites between different samples, we conducted Hierarchical Clustering of all significantly differentiated metabolites and the expression levels of the top 50 differentiated metabolites with VIP values(Figure 6H).The difference compounds were mainly lipids and functional-like molecules (46.7%), followed by phenylpropanoids and polyketides(8.79%) and organic oxygen compounds (8.24%)(Figure 6J). Correlation analysis can help measure the degree of correlation between significantly different metabolites, and further understand the relationship between metabolites in the process of biological state change.The top 50 differential metabolites were selected for visual analysis. Lipids showed positive correlation with lipids and negative correlation with other compounds suggesting that the differential compounds might be regulated by lipids(Figure7C).

**Fig. 7:**
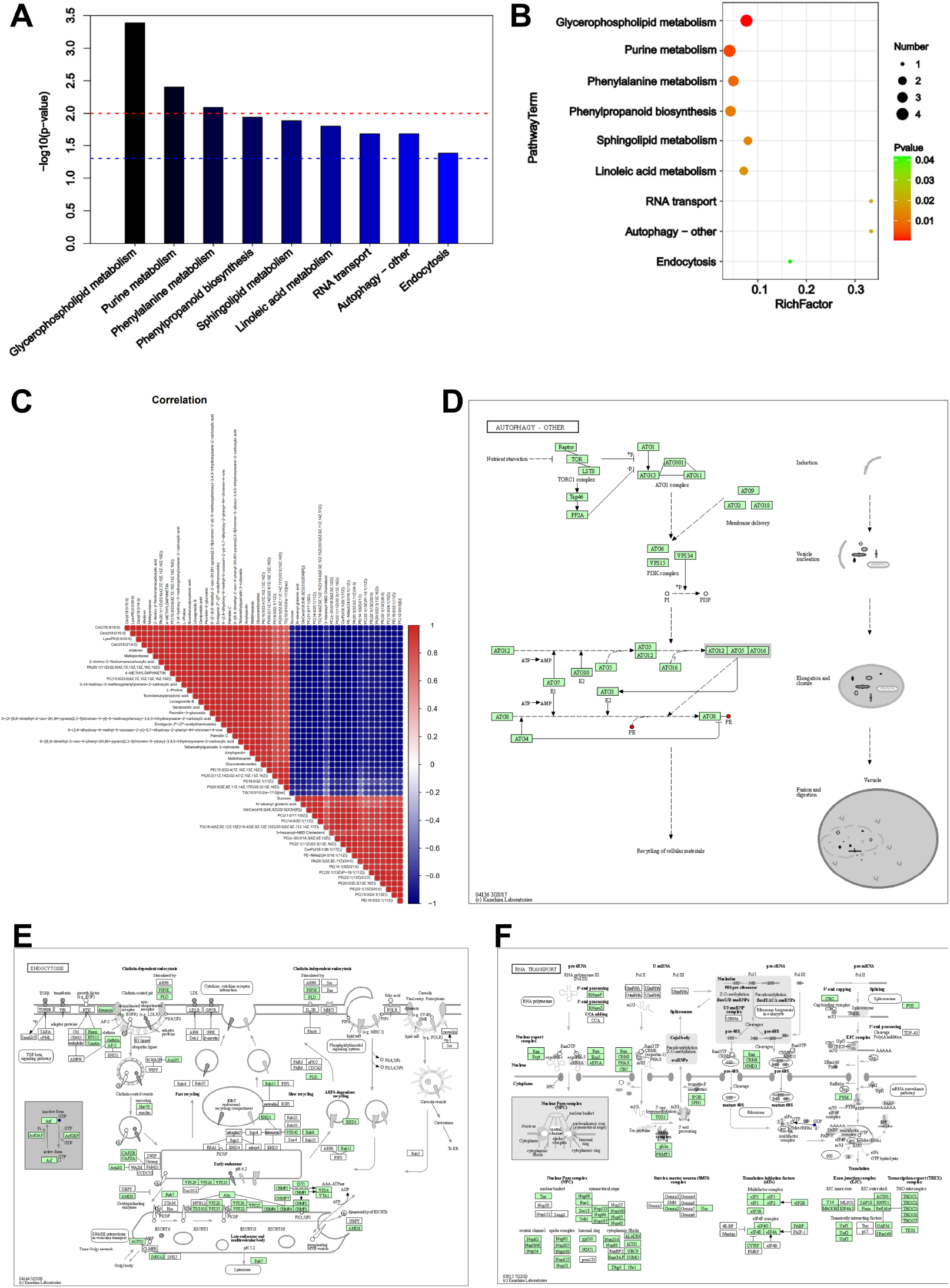
Differential metabolite KEGG enrichment metabolic pathway. (A) Bar graph and (B) Bubble diagram of metabolic pathway enrichment by p-value screening. (C)Correlation analysis of Top 50 differential metabolites.Visualization of (D) Endocytosis, (E) autophagy-other and (F) RNA transport metabolic pathways.

### 3.7 KEGG Analysis

The 182 significantly different metabolites were enriched in 36 pathways, of which a total of 9 pathways were significant (p-value <= 0.05).These were linoleic acid metabolism, RNA transport, autophagy - other, endocytosis, phenylalanine metabolism, phenylpropanoid biosynthesis, sphingolipid metabolism, purine metabolism, and glycerophospholipid metabolism(Figure7A). Further bubble plots of the significantly enriched pathways conducted (Figure7B). The vertical coordinates are the metabolic pathway names and the horizontal coordinates are the enrichment factors.

The differential metabolic pathways are displayed by the KEGG pathway mapper function, and the differential metabolites are colored according to the up- and down-regulation information. The small circles in the metabolic pathway map represent metabolites. Metabolites marked in red in the pathway diagram are experimentally detected up-regulated metabolites and blue are down-regulated metabolites. We visualized 3 metabolic pathway maps for RNA transport, autophagy-other, and endocytosis(Figure7D-F).

## 4 Discussion

Obtaining a large number of plant EVs is still a difficult task due to the limitations of isolation methods. Therefore the study of plant vesicles, especially the function of EVs of medicinal origin, lags far behind the study of mammalian EVs. Several studies have shown that organisms with cell walls such as plants, fungi, gram-positive bacteria and mycobacteria secrete EVs despite being surrounded by a cell wall barrier[28–30]. It is well known that the cell wall is a thicker, tougher structure located outside the cell membrane and is composed of a mucus complex that acts as a protection.The cell wall should prevent anything as large as a EVs passing through it. Although there are some possible resolutions, such as swelling pressure that may force EVs through the cell wall, organisms can release EVs by regulating cell wall thickness, pore size or integrity, or EVs stimulate cell wall remodeling to release it, these approaches are very energy consuming and not easy to implement. Therefore, we believe that studies on EVs extraction from plant apoplastic fluid can only extract a fraction of EVs.Plant EVs are released extracellularly by fusion of multivesicular bodies (MVBs) with the plasma membrane[31], where a small fraction of extracellular vesicles can pass due to the barrier effect of the cell wall. Therefore, we thought of degrading the cell wall using the method of preparing phytoplasma, but with some modifications, and we extracted MOEVs by this method.For comparison, we extracted MONVs using the grinding method. particles larger than 150 nm are readily absorbed by the liver and spleen and are considered unlikely to be used for targeted therapy[32–34]. Therefore, we used 0.22μm membranes and 10,000 g high-speed centrifugation to remove large vesicles during the isolation and purification process.

In this study, we established a method for EVs extraction by enzymatic degradation of plant cell walls, which was validated from the extraction of EVs from MO roots. The contamination of intracellular vesicles was avoided as much as possible during the experiment, such as repeated washing after plant tissue cutting, addition of protoplasm protector and shortening the enzymatic digestion time as much as possible. TEM revealed that the extracted MOEVs had a very similar morphology and structure to mammalian exosomes, were largely intact, and had a uniform and small diameter (50-80nm) (Figure2C and E), similar to the results of a study by Liu et al[35]. The multivariate statistical analysis screened 182 differential metabolites between MOEVs and MONVs, which were mainly enriched in metabolic pathways related to the functions of EVs such as RNA transport, Autophagy - other and Endocytosis.It is not surprising that EVs are enriched in these pathways, as they have been shown to have these functions[31, 36–41]. Due to the change in the methodology of EVs extraction, we extracted EVs with higher yield and purity compared to NVs extracted by conventional juicing.

MOEVs differ from MONVs in terms of lipids, RNA and proteins. In comparison, MOEVs contained less RNA and more protein species and content. Non-targeted metabolomics analysis showed that both vesicles had similar compound class and amounts, but the relative amounts of compound class differed significantly.Both MOEVs and MONVs contained high amounts of lipids (87.62% vs 66.17%), mainly PC, PE, PS, PI, PG and PA, with MOEVs having a higher content of lipids and lipid-like molecules(Figue5F) and smaller size(Figure5C), which may make it more easier to cross the cell wall by deformation. The lipids of plant EVs may themselves act as signaling molecules that accumulate in response to pathogen infection, stress, and play an important role in immunomodulatory defense[36].There are also large differences in lipid composition between MOEVs and MONVs.MOEVs have the most PC (46.40%), followed by PE (29.06%) and PS (7.71%).MONVs have the most PC (29.18%), followed by PE (14.86%) and PI (8.53%). These differences suggest that MONVs are more complex than MOEVs, and we speculate that they are closely related to their origin from mixed particles.

MOEVs and MONVs have different particle sizes, proteins, RNAs and lipids, and there are also different functional differences between MOEVs and MONVs in promoting miR-155 expression in ECs.Recent studies have highlighted the important role of blood vessels in osteogenesis[42, 43], and an increasing number of studies have reported that blood vessels are closely associated with osteogenesis in aged and de-ovulated mice with osteoporosis, where the number of H-type vessels is significantly reduced[1, 44–46]. miR-155 regulates neovascularization and osteogenic differentiation. Recent studies have shown that ECs can also secrete exosomes that specifically target bone tissue and improve osteoporosis by delivering miR-155 in vitro and in vivo[47]. Therefore, promoting high miR-155 expression in ECs is considered as an effective therapeutic strategy to improve osteoporosis. Our study showed that MOEVs have the ability to slightly inhibit miR-155 expression in endothelial cells in a negative correlation at 24h, while substantially promoting miR-155 expression in endothelial cells in a concentration-dependent manner at 48h, for which we do not have more experimental arguments to determine the exact mechanism. Our study also showed that 0.5×10^9^ concentration of MONVs had a strong ability to promote miR-155 expression in endothelial cells at the 48h, which was 2.5-fold higher than the same concentration of MOEVs, although it did not show a stronger effect at a higher concentration of 4.5×10^9^. Apparently MOEVs are also a subgroup of MONVs, but MOEVs in mixed particles could not resolve this phenomenon. There is clearly another type of particle in our experiments that is distinct from MOEVs, and this particle is likely to be an intracellular vesicle that may have better biological activity at lower and safer concentrations, and although methods to isolate subpopulations of particles are still limited, this still raises extensive interest in their subpopulations. These data suggest that we cannot ignore the possibility of intracellular vesicles as therapeutically active agents and drug carriers. Protoplasts prepared by enzymatic cell wall digestion can be further used to study intracellular vesicles.

KEGG analysis of the differential metabolites showed that MOEVs and MONVs have different biological functions, in addition to RNA transport, autophagy (autophagy - other) and endocytosis, the main manifestations of which are linoleic acid metabolism, phenylalanine metabolism, phenylpropanoid biosynthesis, sphingolipid metabolism and purine metabolism.These important differences in biological pathways suggest that these two particles play different biological roles in the growth and development of MO and its environment.

It is important to note that because we applied the enzymatic degradation of cell walls for the extraction of protoplasts very smoothly to the isolation and extraction of plant-derived EVs, we did not explore more about the optimal experimental conditions. Due to the differences in plant species and tissue types, this method must be adapted to the specific plant species and tissue type by appropriately adjusting the type, ratio and time of digestive enzymes, and adjusting the appropriate pH value and adding cytoprotectants is also necessary.

## 5 Conclusion

Our results indicate that degradation of the cell wall could be used to extract large amounts of EVs, a novel method for EVs extraction from plants, and the results also suggest that MOEVs are candidates for novel natural active substances as well as drug carriers that could be elevated by promoting miR-155 expression in endothelial cells. The results confirm that MONVs and MOEVs are distinct particles with differences in size, biological activity and contents. We propose that MOEVs will be used as active substances or drug carriers for the treatment of orthopedic diseases.Although there are some drawbacks in this study, no further animal experiments were conducted to verify its biological activity and toxicity. Due to the advantages of high yield, easy standardization and enzyme recovery, we predict that the use of enzymatic cell wall degradation will be a common method for the preparation of a large number of plant extracellular vesicles.

## CRediT authorship contribution statement

**Qing Zhao**: Design the research, conduct the experiments, and analyze the data and write the original draft. **Guilong Liu**: Designed the research, conducted experiments, and analyzed data, writing original draft. **Manlin Xie**: Conducted experiments and analyzed data. **Yanfang Zou**: Conducted experiments and analyzed data. **Zhaodi Guo**: Conducted experiments and analyzed data. **Fubin Liu**: Conducted experiments and analyzed data. **Jiaming Dong**: Conducted experiments. **Jiali Ye**: Conducted experiments. **Yue Cao**: Analyzed data. **Ge Sun**: Analyzed data. **Lei Zheng**: Supervision, Methodology, Investigation. **Kewei Zhao**: Funding acquisition, Resources, Project administration,Supervision, Methodology, Investigation.

## Acknowledgements

This work was supported by the National Natural Science Foundation of China [grant numbers 81973633], National Natural Science Foundation of China [grant numbers 82174119],Key Laboratory Construction Project of Guangzhou Science and Technology Bureau[grant numbers 202102100007], Collaborative innovation team project of double first-class and high-level universities in Guangzhou University of traditional Chinese Medicine[grant numbers 2021xk62], Youth Innovative Talents Project of Guangdong Province(grant numbers 2020KQNCX015),and Medical Science and Technology Research Fund project of Guangdong Province(grant numbers A2021339).

## A. My Appendix

None

